# SARS-CoV-2-specific T cells associate with reduced lung function and inflammation in pulmonary post-acute sequalae of SARS-CoV-2

**DOI:** 10.1101/2022.02.14.480317

**Authors:** Katherine M. Littlefield, Renée O. Watson, Jennifer M. Schneider, Charles P. Neff, Eiko Yamada, Min Zhang, Thomas B. Campbell, Michael T. Falta, Sarah E. Jolley, Andrew P. Fontenot, Brent E. Palmer

## Abstract

As of January 2022, at least 60 million individuals are estimated to develop post-acute sequelae of SARS-CoV-2 (PASC) after infection with severe acute respiratory syndrome coronavirus 2 (SARS-CoV-2). While elevated levels of SARS-CoV-2-specific T cells have been observed in non-specific PASC, little is known about their impact on pulmonary function which is compromised in the majority of these individuals. This study compares frequencies of SARS-CoV-2-specific T cells and inflammatory markers with lung function in participants with pulmonary PASC and resolved COVID-19 (RC). Compared to RC, participants with respiratory PASC had up to 34-fold higher frequencies of IFN-γ- and TNF-α-producing SARS-CoV-2-specific CD4^+^ and CD8^+^ T cells in peripheral blood and elevated levels of plasma CRP and IL-6. Importantly, in PASC participants the frequency of TNF-α-producing SARS-CoV-2-specific CD4^+^ and CD8^+^ T cells, which exhibited the highest levels of Ki67 indicating they were activity dividing, correlated positively with plasma IL-6 and negatively with measures of lung function, including forced expiratory volume in one second (FEV1), while increased frequencies of IFN-γ-producing SARS-CoV-2-specific T cells associated with prolonged dyspnea. Statistical analyses stratified by age, number of comorbidities and hospitalization status demonstrated that none of these factors affect differences in the frequency of SARS-CoV-2 T cells and plasma IL-6 levels measured between PASC and RC cohorts. Taken together, these findings demonstrate elevated frequencies of SARS-CoV-2-specific T cells in individuals with pulmonary PASC are associated with increased systemic inflammation and decreased lung function, suggesting that SARS-CoV-2-specific T cells contribute to lingering pulmonary symptoms. These findings also provide mechanistic insight on the pathophysiology of PASC that can inform development of potential treatments to reduce symptom burden.

**Author Summary:** Long COVID-19 or post-acute sequelae of SARS-CoV-2 (PASC) impacts 20-30% of those infected with SARS-CoV-2 and is characterized by COVID-19 symptoms exceeding 4 weeks from symptom onset. While those with PASC experience a wide variety of persistent symptoms including shortness of breath, cough, chest pain, irregular heartbeat, brain fog, fatigue, and intermittent fever, lung-related conditions are the most common. Although, infection with SARS-CoV-2 is clearly the inciting factor for PASC, the mechanisms responsible for long-term lung dysfunction are unclear and current treatments are ineffective at resolving pulmonary symptoms. Generalized PASC has been associated with SARS-CoV-2-specific T cells, a component of adaptive immunity, suggesting that residual virus may persist. Here, we investigated the frequency and function of virus-specific T cells in the blood of individuals with pulmonary PASC and correlated their presence with systemic inflammation and lung function. Our findings demonstrated that T cells specific for SARS-CoV-2 are elevated in the blood of those with pulmonary PASC and are associated with increased IL-6, a cytokine strongly associated with COVID-19 severity, and decreased lung function. These findings provide mechanistic insight into the pathophysiology of pulmonary PASC needed for the development of new treatments to improve quality of life for those affected.

## Introduction

After infection with severe acute respiratory syndrome coronavirus 2 (SARS-CoV-2), 20-30% of survivors experience prolonged symptoms that can significantly impact quality of life(1). “Long-COVID” or “Long-haul COVID” refers to individuals experiencing persistent symptoms that can involve multiple organ systems, including the lungs, heart, and brain(2–4). Officially named post-acute sequelae of SARS-CoV-2 (PASC), this syndrome is defined as new, continuing or recurring symptoms of COVID-19 that occur four or more weeks after initial infection(1). Hallmark symptoms of PASC include persistent palpitations, neuropsychiatric conditions, anosmia and dysgeusia, with dyspnea and other respiratory ailments being the most common(5–7). Reduced lung volume and exercise capacity are commonly observed in survivors of COVID-19 pneumonia(8), however, the appearance and persistence of PASC respiratory symptoms is not related to the severity of initial illness(9). As new SARS-CoV-2 variants potentially increase infection rates and disease severity(10, 11), mutations to viral surface proteins may also increase the prevalence of persistent symptoms(12). Early reports show sustained frequencies of SARS-CoV-2-specific T cells and elevated inflammatory systemic markers have been observed in non-specific PASC(13, 14), and understanding the immunologic mechanisms of pulmonary PASC is of vital importance for developing treatment options to reduce symptom burden.

The T cell adaptive immune response is well characterized in acute and convalescent cases of COVID-19 and contributes to virus clearance, protective immunity and inflammation(15). The frequency of SARS-CoV-2-specific T cells positively correlates with both serum antibody levels and disease severity; however, while SARS-CoV-2-specific antibodies remain relatively stable up to 240 days, virus-specific CD4^+^ and CD8^+^ T cell frequencies decline with a half-life of 3-5 months(16, 17). In mild/asymptomatic cases, SARS-CoV-2-specific T cells are polyfunctional and produce multiple cytokines(18); conversely, during severe disease, polyfunctional virus-specific T cells are underrepresented and are skewed towards a cytotoxic phenotype(19). Although SARS-CoV-2-specific T cells are protective in most cases, it has been shown they can contribute to the cytokine release syndrome seen in patients with severe COVID-19(20). Furthermore, CD8^+^ T cells in the lung during acute infection are associated with inflammation, fibrosis, biomarkers of vascular injury, and poor outcomes(21, 22). Thus, while SARS-CoV-2-specific T cells likely play a role in PASC, the characteristics of these cells and their connections to systemic inflammation or pulmonary symptoms are currently unknown.

Here we determined the frequency and function of SARS-CoV-2-specific T cells in blood, and their relationship with the expression of plasma inflammatory markers and measures of lung function in individuals with pulmonary PASC. We found patients with pulmonary PASC had significantly elevated frequencies of IFN-γ- and TNF-α-producing SARS-CoV-2-specific T cells compared to participants with resolved COVID-19 (RC). These virus-specific T cells were strongly associated with increased markers of inflammation and decreased lung function in PASC. These findings indicate pulmonary PASC may be, in part, driven by the production of inflammatory cytokines by SARS-CoV-2-specific T cells.

## Results

### Cohort Descriptions

Study participants were recruited between July 2020 and April 2021, prior to appearance of the B.1.617.2 or B.1.1.529 variants in Colorado(23). Patients were confirmed SARS-CoV-2 PCR positive by nasopharyngeal swab during the acute phase of infection. Participants categorized as pulmonary PASC experienced prolonged tussis, dyspnea and/or fatigue (S1 Table). The pulmonary PASC cohort reported symptoms lasting for a median duration of six months from symptom onset or hospital discharge. All RC participants reported no symptoms at the time of sample collection, and if RC participants subsequently did experience relapse of symptoms, they were excluded from the study. All participants with chronic or active infections other than SARS-CoV-2, using medications targeting IL-6, or antibiotic use within one month of sample collection were also excluded.

Clinical characteristics of the PASC and RC cohorts were similar to those observed by other groups(1, 9) and are highlighted in Table 1. Those with PASC were older than RC participants (median years (range), PASC=54 (22–69), RC=33 (22–71), P=0.003) and 40% required hospitalization (duration 3-52 days: median=11 (P=0.01)) during acute COVID-19 infection (Table 1). Overall, no significant differences between PASC and RC cohorts in terms of pre-existing conditions were found (Table 1). Those with PASC experienced an average symptom duration of over 6 months while RC participants’ average symptom duration was 12 days (P<0.0001) (Fig. 1a). PASC participants reported a median of 9 symptoms while RC participants reported 6 symptoms during initial infection (P=0.002). The median number of prolonged symptoms reported by those with PASC was 5 while those with RC reported none (P<0.0001) (Fig. 1b). There were no significant differences in the total duration of symptoms between PASC participants who were hospitalized (PASC-H) and those with PASC who were not hospitalized (PASC-NH) (P=0.17). PASC-NH participants reported a greater variety of symptoms during both the acute (P=0.03) and post-acute phases (P=0.02) of disease when compared to symptoms reported by PASC-H participants (Fig 1b). The average time from symptom onset to blood collection was 225 days for the PASC and 32 days for the RC cohorts (Table 1). Statistical analyses stratified by age (S1 and S4 Fig.), number of comorbidities (S1 and S4 Fig.), time to sample collection (S2 Fig.), and hospitalization status (Fig. 6a) demonstrated that none of these factors affected the differences in frequency of SARS-CoV-2 T cells and plasma IL-6 levels measured between PASC and RC cohorts.

**Fig 1.**
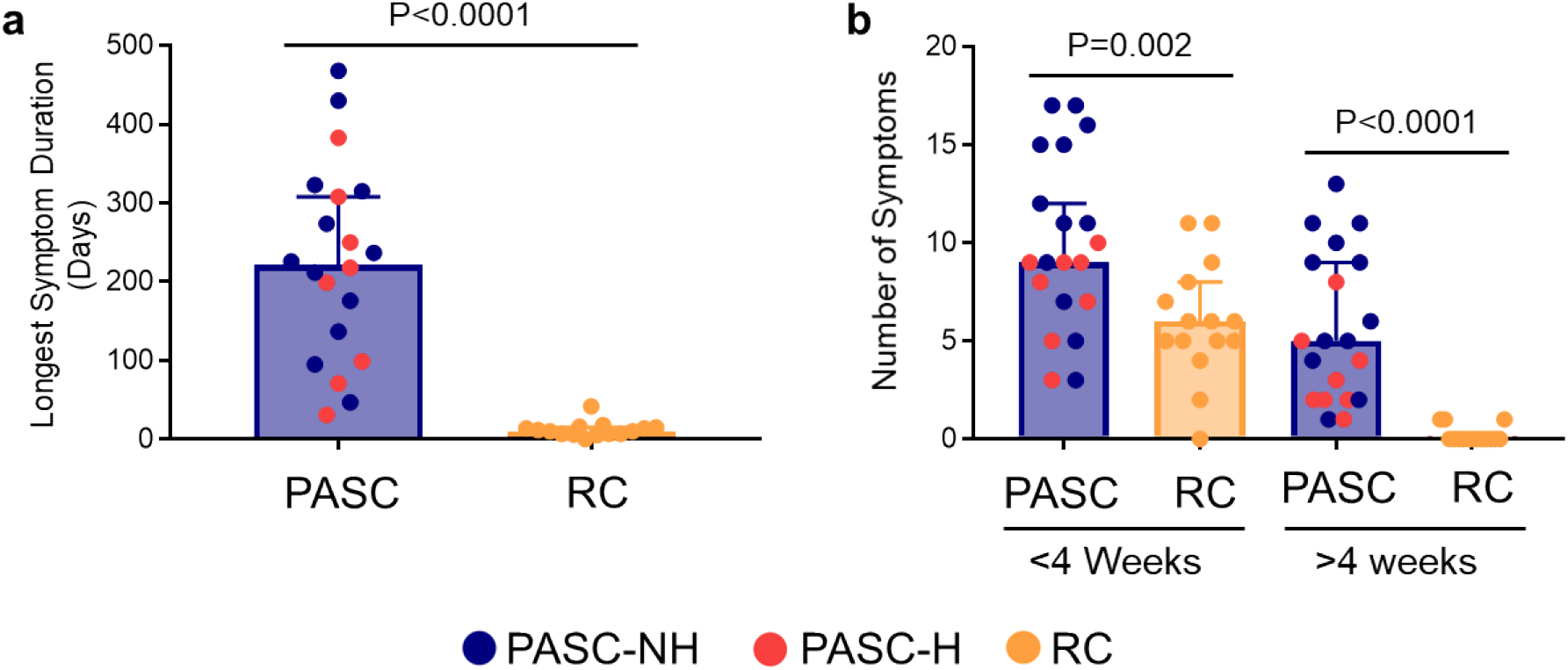
Symptom characteristics of PASC and RC participants. (a) Symptom duration (days) reported in symptom questionnaires for PASC and RC participants. (b) Number of symptoms reported <4 weeks or >4 weeks from symptom onset for PASC and RC participants. For each graph, the horizontal bars represent the median of each cohort and the error bars represent the 95% confidence interval. Blue represents PASC participants not hospitalized (PASC-NH, n=12), red represents PASC-hospitalized (PASC-H, n=8) and orange represents RC participants (n=15). Mann-Whitney tests were used to determine statistical significance.

**Table 1.** Demographics of Cohorts.

|  | Cohort |  | P Value<sup>a</sup> | PASC Hospitalized |  | P Value<sup>a</sup> |
| --- | --- | --- | --- | --- | --- | --- |
|  | PASC | RC |  | Yes | No |  |
| Number of participants<sup>b</sup> | 20 | 15 |  | 8<sup>Δ</sup> | 12 |  |
| Female | 10 (50) | 5 (33) | ns | 2 (25) | 8 (75) | ns |
| Male | 10 (50) | 10 (66) | ns | 6 (75) | 4 (25) | ns |
| Median time to sample<sup>c</sup><br>(Range) | 212 (64-396) | 32 (13-265) | ***<sup>Δ</sup> | 184.5 (31-383) | 183 (45-332) | ns |
| Median Age (Range) | 53 (22-65) | 34* (22-71) | **<sup>Δ</sup> | 60 (49-69) | 50 (22-62) |  |
| ICU Admission | 6 (32) | NA<sup>d</sup> | NA | 6 (75) | NA | NA |
| **Race and Ethnicity** |  |  |  |  |  |  |
| White | 17 | 14 | ns | 6 | 11 | ns |
| Black | 2 | 1 | ns | 1 | 1 | ns |
| American Indian/Alaska<br>Native | 0 | 2 | ns | 0 | 0 | ns |
| Other | 1 | 0 | ns | 1 | 0 | ns |
| Hispanic or Latin Origin | 5 | 4 | ns | 3 | 2 | ns |
| **Underlying Medical Condition** |  |  |  |  |  |  |
| Any | 10 (50) | 3 (20) | ns<sup>Δ</sup> | 4 (50) | 6 (50) | ns |
| Hypertension | 7 (35) | 0 (0) | * | 2 (38) | 4 (33) | ns |
| Pulmonary Disease | 2 (10) | 1 (7) | ns | 0 (0) | 2 (17) | ns |
| Immune System Disease | 1 (5) | 0 (0) | ns | 0 (0) | 1 (8) | ns |
| Cancer | 1 (5) | 0 (0) | ns | 0 (0) | 1 (8) | ns |
| Kidney Disease | 1 (5) | 1 (7) | ns | 0 (0) | 1 (8) | ns |
| Metabolic Disease | 5 (25) | 0 (0) | ns | 4 (50) | 1 (8) | ns |
| **Medications during Hospitalization** |  |  |  |  |  |  |
| Convalescent Plasma | 3 (15) | NA | NA | 3 (38) | NA | NA |
| Hydroxychloroquine | 4 (20) | NA | NA | 4 (50) | NA | NA |
| Remdesivir | 4 (20) | NA | NA | 4 (50) | NA | NA |
| Dexamethasone | 4 (20) | NA | NA | 4 (50) | NA | NA |
| Tocilizumab | 0 (0) | NA | NA | 0 (0) | NA | NA |

<sup>a</sup>Mann-Whitney tests were used to determine statistical significance, ns = not significant \* = P<0.05, \*\* = P<0.005, \*\*\* = P<0.0005.
<sup>b</sup>All values are number of participants with the percentage of the cohort in parentheses unless otherwise specified.
<sup>c</sup>Median number of days from first reported symptom to collection of blood.
<sup>d</sup>NA=Not applicable.
<sup>Δ</sup>Indicates stratification analyses were performed demonstrating these differences do not influence SARS-CoV-2-specific T cell frequencies (S1 and S2 Fig.) and levels of plasma IL-6 (S4 Fig.).

### Elevated SARS-CoV-2-specific T cells in pulmonary PASC

We measured the frequency of SARS-CoV-2-specific T cells in blood using intracellular cytokine (IFN-γ, TNF-α, and IL-2) staining after stimulation with peptide pools of the SARS-CoV-2 spike (S), nucleocapsid (N) or membrane (M) surface-expressed proteins. Representative density plots of SARS-CoV-2-specific T cell populations are shown (Fig. 2a). First, we analyzed the combined frequency of SARS-CoV-2-specific T cells for all three proteins. PASC participants had significantly increased frequencies of SARS-CoV-2-specific CD4^+^ and CD8^+^ T cells that produced IFN-γ or TNF-α compared to the RC cohort, while frequencies of IL-2-producing CD4^+^ SARS-CoV-2-specific T cells also trended higher (Fig. 2b-g). There was a 2.9- and 5.2-fold increased frequency of CD4^+^ and CD8^+^ SARS-CoV-2-specific T cells producing IFN-γ in PASC participants compared to RC participants (Fig. 2b-c). The differences in the frequencies of TNF-α-producing CD4^+^ and CD8^+^ T cells in PASC participants compared to RC participants were even greater (6.9- and 34-fold, respectively) (Fig. 2d-e). We chose to separate pulmonary PASC participants by prior hospitalization status to determine if this factor was associated with the frequency of virus-specific T cells. The same significant differences were observed when comparing only PASC-NH and RC participants and no significant differences were observed between PASC-NH and PASC-H groups (Fig. 2b-g). Stratification analyses for age, comorbidities and time of sample collection showed no significant differences within the RC or pulmonary PASC cohorts (S1 and S2 Fig).

**Fig 2.**
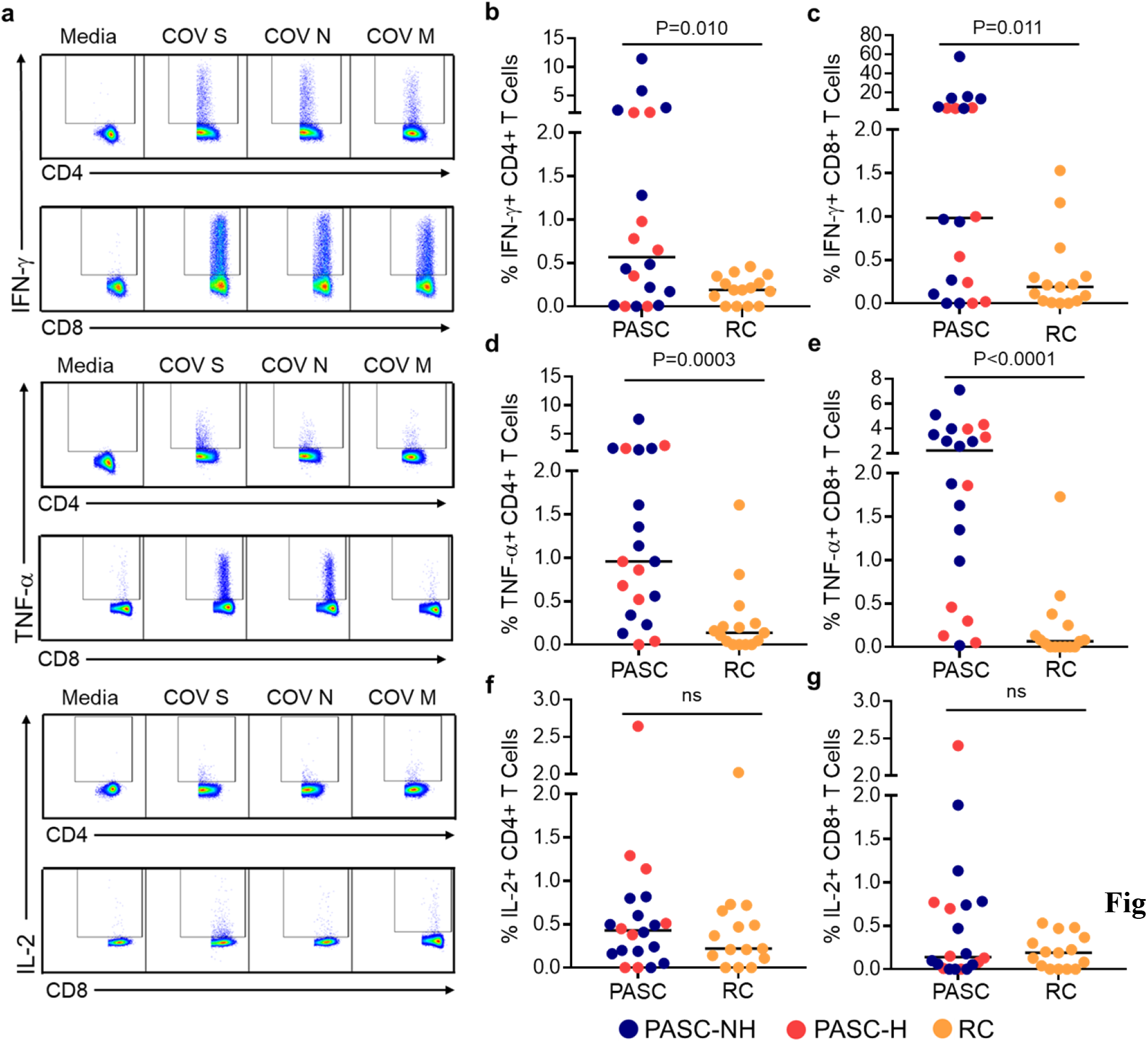
Cumulative SARS-CoV-2 specific T cell frequencies are elevated in PASC. (a) Representative flow cytometry density plots of SARS-CoV-2-specific T cells stimulated with S, N and M peptide pools (5 μg/ml) for six hours with from one participant with PASC. Samples were gated through lymphocytes, live, CD3+, separated by CD4+/CD8+ and then frequencies of cytokines were assessed. Percent of total CD4^+^ producing (b) IFN-γ, (c) TNF-α, (d) IL-2 or CD8^+^producing (e) IFN-γ, (f) TNF-α or (g) IL-2 T cells in response to S, N and M peptide pools. Each point represents the sum of the combined frequencies of virus-specific T cells to the peptide pools from each participant. The horizontal bars depict the median values for each cohort. Color labeling the same as in Figure 1. Mann-Whitney tests were used to determine statistical significance.

Next, we individually assessed IFN-γ- and TNF-α-producing SARS-CoV-2-specific T cell frequencies as these cytokines had the most significant differences overall. The frequency of IFN-γ-producing, S-specific CD4^+^ T cells was significantly higher in those with pulmonary PASC as compared to RC participants (median, range; PASC: 0.23%, 0-7.63%; RC: 0.075%, 0-0.26%: P=0.0098) (Fig. 3a). No significant differences were noted in the frequency of SARS-CoV-2 N- and M-specific CD4^+^ T cells producing IFN-γ in PASC and RC participants (Fig. 3a). Similar findings were seen for IFN-γ expression in CD8^+^ T cells between PASC and RC cohorts and again no difference was observed comparing the PASC-NH and PASC-H groups (Fig. 3b). Again, there were no significant differences based on age, time of sample collection or comorbidities for IFN-γ-producing SARS-CoV-2 (S1 and S2 Fig).

**Fig 3.**
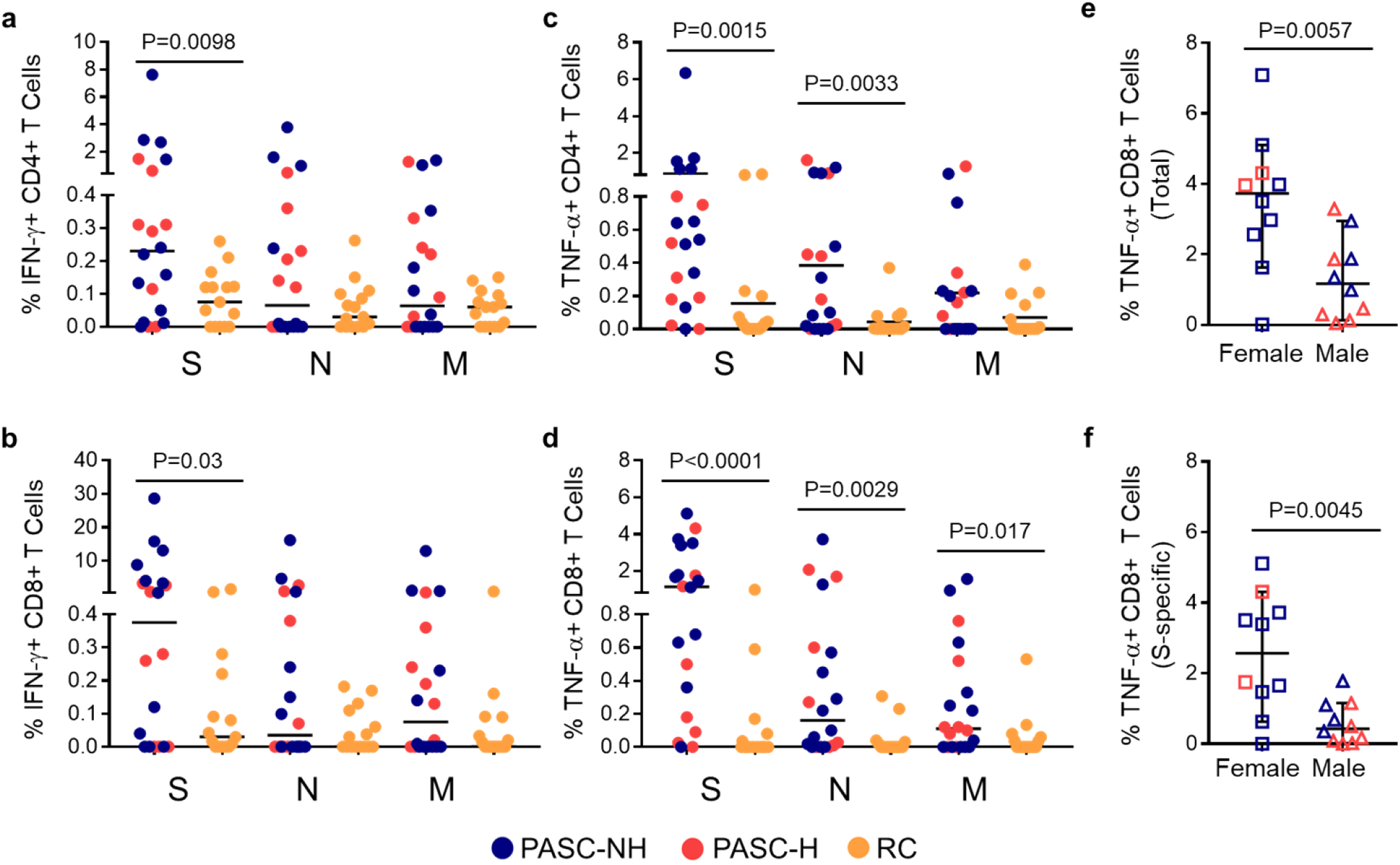
IFN-γ and TNF-α producing T cell responses to individual SARS-CoV-2 proteins. Percent of CD4^+^ T cells producing (a) IFN-γ or (b) TNF-α and CD8^+^ T cells producing (c) IFN-γ or (d) TNF-α in response to S, N and M peptide pools separately. (e) Frequency of total (S, N, and M) CD8^+^ T cells or (f) S-specific CD8^+^ T cells producing TNF-α for PASC participants compared by sex. Blue represents PASC-NH (not hospitalized), red represents PASC-hospitalized and orange represents RC participants. Mann-Whitney tests were used to determine statistical significance.

PASC participants had significantly higher frequencies of TNF-α-producing SARS-CoV-2 S- and N-specific CD4^+^ T cells, (P=0.0015 and P=0.0033, respectively) (Fig. 3c) and significantly increased frequencies of TNF-α-producing CD8^+^ T cells in response to all three SARS-CoV-2 proteins (Fig. 3d) compared to RC participants. Approximately 50% of CD4^+^ and CD8^+^ T cells from PASC participants produced TNF-α in response to all 3 SARS-CoV-2 proteins, whereas these percentages were 33% and 13%, respectively in RC participants. Only one PASC participant had no detectable CD4^+^ T cell cytokine response to any SARS-CoV-2 protein – this individual did have CD8^+^ T cell cytokine responses – while 5 RC participants had no detectable responses in either CD4^+^ or CD8^+^ T cells. Interestingly, female PASC participants had significantly higher total (P=0.0015) and S-specific (P=0.045) CD8^+^ T cell responses compared to male participants with PASC (Fig. 3e-f).

### SARS-CoV-2-specific T cells in PASC are less polyfunctional than in RC and exhibit recent proliferation

Next, we compared the expression of cytokines and phenotypic markers on SARS-CoV-2-specific T cells in pulmonary PASC and RC participants. It has been established that T cell immunity to SARS-CoV-2 wanes rapidly after resolution of infection and symptoms(24). As confirmation of waning immunity, we examined SARS-CoV-2-specific T cell frequencies in 2 RC participants who provided blood at 2- and 30-weeks post resolution of symptoms. As expected, their T cell responses decreased over time (S2 Fig). Based on these data, we collected blood from RC participants at early time points after resolution of infection to ensure that detectable frequencies of SARS-CoV-2-specific T cells were present. Thus, we were able to interrogate differences in T cell phenotype and function in both PASC and RC participants.

We assessed the cytokine-production profiles of PASC and RC participants utilizing simplified presentation of incredibly complex evaluations (SPICE)(25). The SPICE analysis revealed the majority of SARS-CoV-2-specific T cells in individuals with pulmonary PASC only produce one of the three cytokines tested with TNF-α dominating the virus-specific CD4^+^ T cell response and IFN-γ dominating the CD8^+^ T cell response, while RC participants tended to produce multiple cytokines, indicating the T cell cytokine response is more restricted PASC compared to RC. A deeper analysis revealed significant differences in the distribution of CD4^+^ T cell cytokine expression in response to N and M proteins when comparing PASC and RC participants (P=0.012 and P=0.046, respectively) (Fig. 4). The proportion of N-specific CD4^+^ T cells producing both IFN-γ and IL-2 was significantly higher in RC participants compared to PASC participants (P=0.017) while the proportion of TNF-α- and IL-2-producing N-specific CD4^+^ T cells was higher in pulmonary PASC participants (P=0.024). For CD8^+^ T cells, the overall proportions of cytokine co-expression were also significantly different between PASC and RC cohorts for S- and N-specific T cells (P=0.012 and P=0.008, respectively) and although at an overall low frequency, the proportion of TNF-α- and IL-2-producing N-specific CD8^+^ T cells was also higher in PASC compared to RC participants (Fig. 4). Interestingly, the proportion of IFN-γ- and TNF-α-producing S-specific CD8^+^ T cells was significantly greater in RC compared to PASC (P=0.026), while the proportion of CD8^+^ T cells secreting this combination of cytokines trended higher in PASC in response to N- and M-peptide pools.

**Fig 4.**
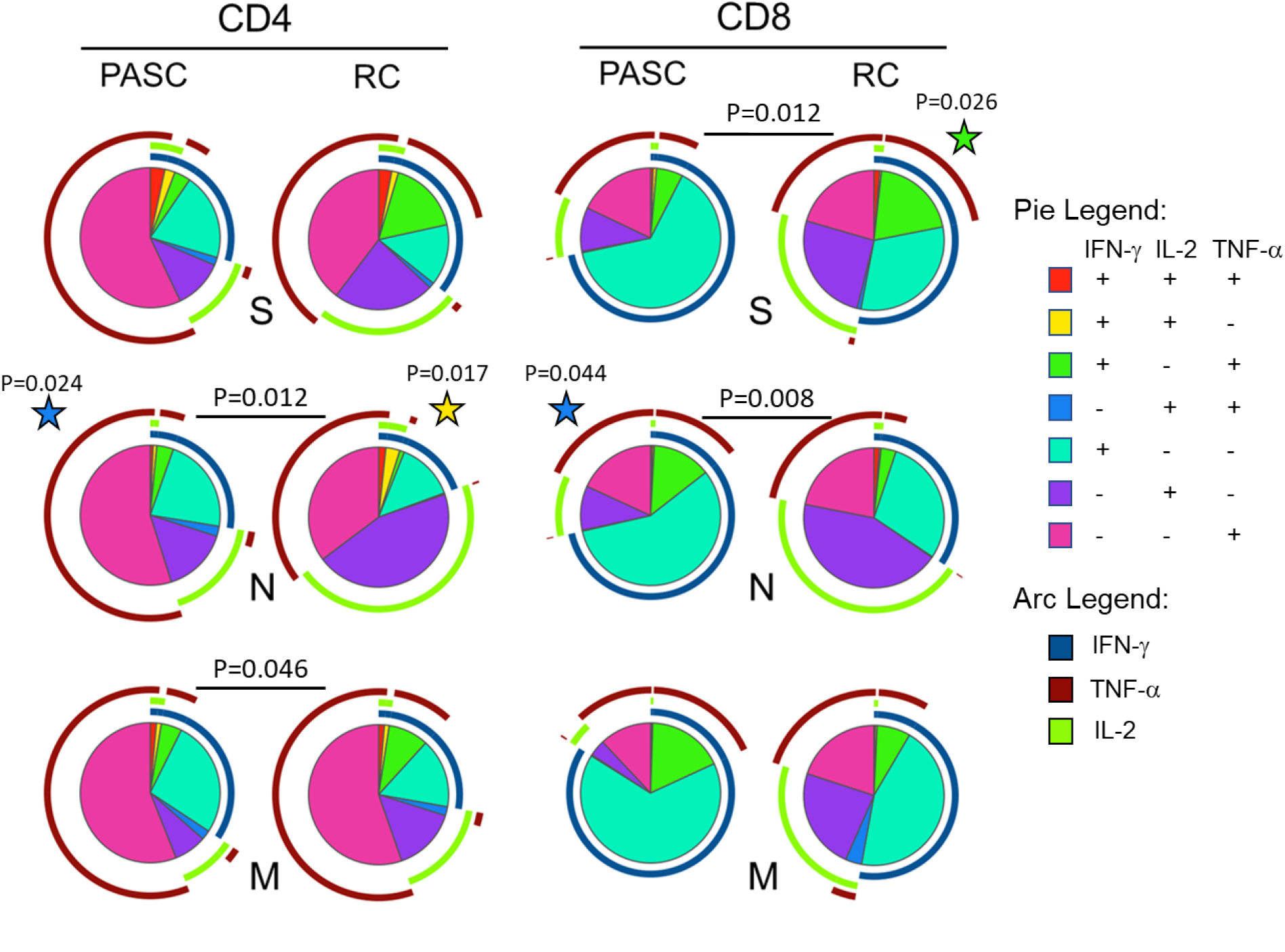
Cytokine co-expression of SARS-CoV-2 specific T cells differs between PASC and RC. Cytokine co-expression on SARS-CoV-2 specific T cells visualized using simplified presentation of incredibly complex evaluations (SPICE) analysis. Each pie chart represents the proportions of combinations of IFN-γ, TNF-α and IL-2 producing T cells in response to one SARS-CoV-2 protein. Arcs surrounding each pie chart depict the proportion of cells secreting each individual cytokine. Colors for pie charts and arcs represent different cytokines or combinations of cytokines and are listed in their corresponding legend. Stars denote significant differences determined by student t test between PASC and RC cohorts for a particular combination of co-expressed cytokines matching as indicated by the color corresponding to the pie legend. Stars are positioned next to the cohort with the higher proportion. P values positioned between PASC and RC pie charts denote statistical significance of overall composition by permutation test with 10,000 iterations.

We then assessed markers of T cell maturation (CD27 and CD45RA), exhaustion (PD-1) and proliferative capacity (Ki-67) on total (S3 Fig) and virus-specific (Fig. 5) T cells from PASC and RC participants. No differences were seen in the frequency of naïve, effector memory or terminally differentiated effector memory for total CD4^+^ or CD8^+^ T cells between the two groups; however, there was an increased frequency of central memory (CD27^+^ CD45RA^-^) CD4^+^ T cells in the blood of PASC participants (PASC median: 43%, RC median: 35%; P=0.04) (S3 Fig). Also, no significant differences in Ki-67 or PD-1 expression were seen. Regarding SARS-CoV-2-specific T cells, maturation and exhaustion markers were not significantly different between PASC and RC participants (data not shown). However, within the pulmonary PASC cohort, Ki-67 expression in SARS-CoV-2-specific TNF-α-producing CD4^+^ and CD8^+^ T cells was significantly higher than in cells expressing either IFN-γ or IL-2 (Fig. 5). For example, in response to M protein, the number of TNF-α-producing cells expressing Ki-67 was 2.6-fold higher for CD4^+^ T cells (P<0.0001) and 3.2-fold higher for CD8^+^ T cells (P=0.0059) compared to IFN-γ-producing T cells (Fig. 5). TNF-α-producing T cells exhibited significantly higher frequencies of Ki-67 than IFN-γ-producing T cells for both CD4^+^ and CD8^+^ T cell subsets, and Ki-67 was higher on TNF-α-producing T cells for all conditions, except for S-specific CD4^+^ T cells when compared to the frequency of Ki-67 on IL-2-producing T cells (Fig. 5). For S-specific CD8^+^and N-specific CD4^+^ T cells the frequency of Ki-67 on IL-2-producing T cells was somewhat higher than that of IFN-γ-producing T cells (P=0.03 and P=0.04, respectively) (Fig. 5).

**Fig 5.**
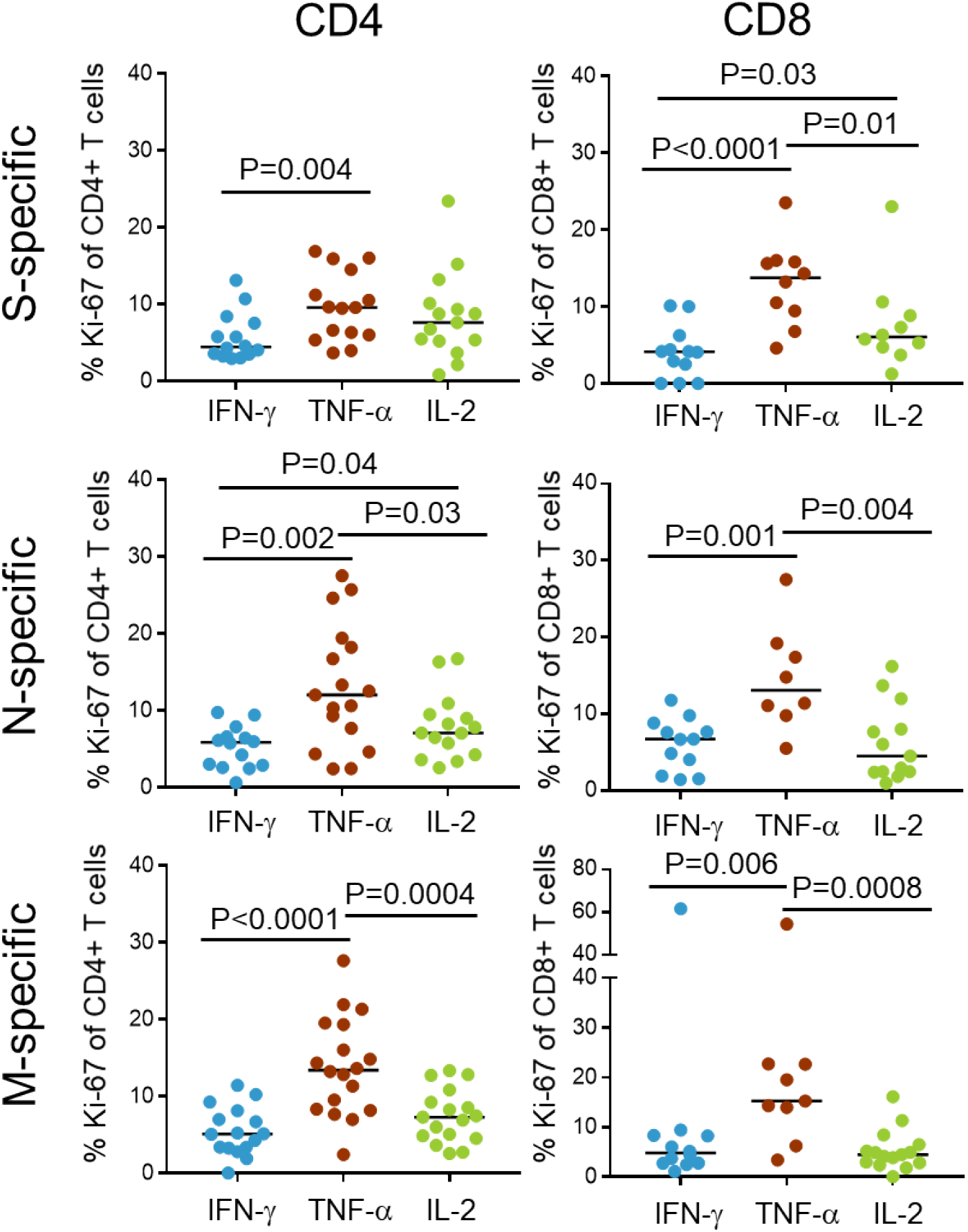
TNF-α-producing T cells have the highest proportion of Ki-67 expression. Shown are the percentages of CD4 T cells (left panels) and CD8 T cells (right panels) obtained from the blood of PASC participants that are positive for Ki-67 expression. T cell populations are further grouped by their production of cytokines and responses to peptide pools of SARS-CoV-2 structural proteins (S, top panels; N, middle panels; M, bottom panels). Note, data points from individual PASC participants were obtained for 1 or more of the cytokines assessed; however, in no instances are there multiple values obtained from the same participant for a particular cytokine. Blue represents IFN-γ^+^ T cells, brown represents TNF-α^+^ T cells and green represents IL-2^+^ T cells. Mann-Whitney tests were used to determine statistical significance.

### Plasma IL-6 levels in pulmonary PASC correlated with the frequency of SARS-CoV-2-specific T cells

We measured plasma IL-6 and CRP in participants to characterize systemic inflammation in pulmonary PASC and correlate these markers with the frequency of virus-specific T cells. Assessed independently of hospitalization status, both IL-6 and CRP were significantly elevated compared to the RC cohort: IL-6 (PASC median=2.9 pg/mL, RC median=1.7 pg/mL, P=0.025); CRP (PASC median=4.4 mg/L, RC median=1.76 mg/L, P=0.0044) (Fig. 6a-b). No significant correlations were found between plasma IL-6 or CRP levels and age or number of pre-existing conditions for all participants or each cohort separately (data not shown). No significant difference in plasma IL-6 between PASC-H and PASC-NH were found and both were significantly elevated compared to RC (data not shown). There was also no difference in IL-6 comparing female and male PASC participants (Fig. 6b). Assessing CRP in PASC-NH participants, this group trended higher than RC participants (PASC-NH median=3.10 mg/L, RC median=1.76 mg/L, P=0.074), whereas there was a significant difference between PASC-H and RC (PASC-H median=5.74 mg/L, RC median=1.76 mg/L, P=0.004). Of note, PASC-H participants were significantly higher compared to PASC-NH (P=0.025) (data not shown). Male PASC participants also had significantly higher plasma CRP compared to female PASC participants (P=0.028) (Fig. 6d). This observation suggests that elevated plasma CRP in pulmonary PASC is likely related to initial disease severity, known to be associated with male sex(26), while IL-6 elevation is specific to pulmonary PASC regardless of disease severity or gender. Stratification analyses show no significant differences within the PASC or RC cohorts based on age or pre-existing conditions, although CRP did trend higher in those with pre-existing conditions (S4 Fig). No correlations between duration of symptoms, time from onset to sample collection or age with IL-6 or CRP were found (data not shown). We also compared IgG and IgA antibody levels to the S1 region of the spike protein and found no differences when comparing all PASC or PASC-NH participants with the RC cohort (IgG: P=0.45, IgA: P=0.43): however, PASC-H participants had significantly higher IgG and IgA antibody levels than PASC-NH participants (IgG: P=0.007, IgA: P=0.007) (S4 Fig).

**Fig 6.**
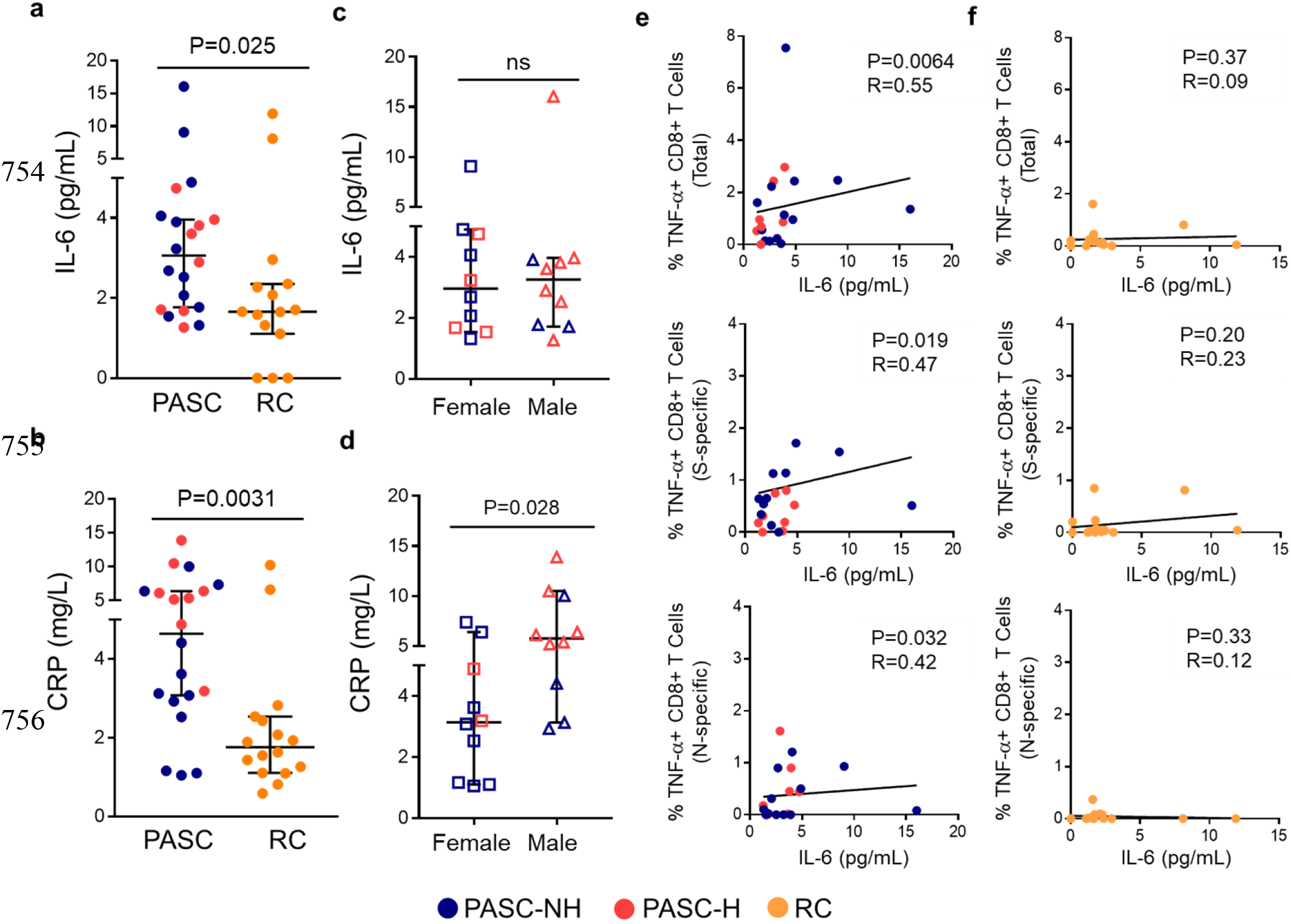
Plasma IL-6 in PASC is higher than RC and correlates with frequency of TNF-α producing CD8^+^ T cells. (a) Plasma IL-6 levels (pg/mL) and (b) plasma CRP levels (mg/L) in PASC versus RC. (c) PASC plasma IL-6 (pg/mL) levels and (d) PASC plasma CRP levels (mg/L) in female versus male. € Correlations between plasma IL-6 and frequency of TNF-α-producing CD8^+^ T cells in PASC participants. (f) Correlations between plasma IL-6 and frequency of TNF-α-producing CD8^+^ T cells in RC participants. For a-d, bar represents median of cohort and error bar is 95% confidence index. Each point represents data from one participant. Blue: PASC-NH (not hospitalized), red: PASC-Hospitalized and orange: RC participants. For a-d Mann-Whitney tests were used to determine statistical significance. For e-f Spearman correlations were used to determine statistical significance.

We next explored the relationship between the frequencies of SARS-CoV-2-specific CD4^+^ and CD8^+^ T cells with IL-6 and CRP. We identified significant positive correlations between total, S-specific, and N-specific frequencies of TNF-α-producing SARS-CoV-2-specific CD8^+^ (r=0.55; P=0.0064, r=0.47; P=0.019, and r=0.42; P=0.032 respectively) T cells and plasma IL-6 in PASC participants (Fig. 6e). These correlations were not observed in the RC cohort (Fig. 6f). No significant correlations between plasma CRP and the frequency of SARS-CoV-2-specific T cells in either PASC or RC cohorts were observed (data not shown).

### SARS-CoV-2-specific T cell frequencies correlate with decreased lung function

A subset of pulmonary PASC participants (n=8) had pulmonary function tests (PFTs) performed during their period of prolonged respiratory symptoms as part of their standard of care. None of these participants reported pre-existing pulmonary conditions prior to infection with SARS-CoV-2. PFTs were performed between 45 and 315 days after symptom onset (median=187 days). We correlated the frequencies of SARS-CoV-2-specific T cells with the following variables: percent predicted forced vital capacity (%FVC), absolute and percent predicted forced expiratory volume during the 1^st^ second (FEV_1_, %FEV_1_ respectively), FEV_1_/FVC, total lung capacity percent predicted (%TLC), single-breath diffusing capacity of the lung for CO percent predicted (%DLCO_SB), and diffusing capacity of the lung per alveolar volume percent predicted (%DLCO/VA). As shown in Fig. 7a and 7b, the total frequencies of IFN-γ-producing SARS-CoV-2-specific CD4^+^ and CD8^+^ T cells negatively correlated with %FEV_1_ (r=-0.81, P=0.011; r=-0.9, P=0.007, respectively). Similar findings were seen between TNF-α-producing CD4^+^ and CD8^+^ T cells and %FEV_1_ (Fig. 7c-d). We then compared the frequency of SARS-CoV-2-specific T cells with the duration of prolonged dyspnea experienced by 80% (n=16) of our pulmonary PASC participants. From this analysis, we identified positive correlations between dyspnea duration and frequencies of IFN-γ-producing SARS-CoV-2 total (r=0.49, P=0.02) and S-specific (r=0.55, P=0.015) CD8^+^ T cells (Fig 7e-f). There was also a positive correlation between TNF-α-producing SARS-CoV-2 S-specific (r=0.45, P=0.036) CD8^+^ T cells and dyspnea duration, and a negative correlation between total CD4^+^ IL-2-producing T cells and dyspnea duration (r=-0.61, P=0.006) (Fig g-h).

**Fig 7.**
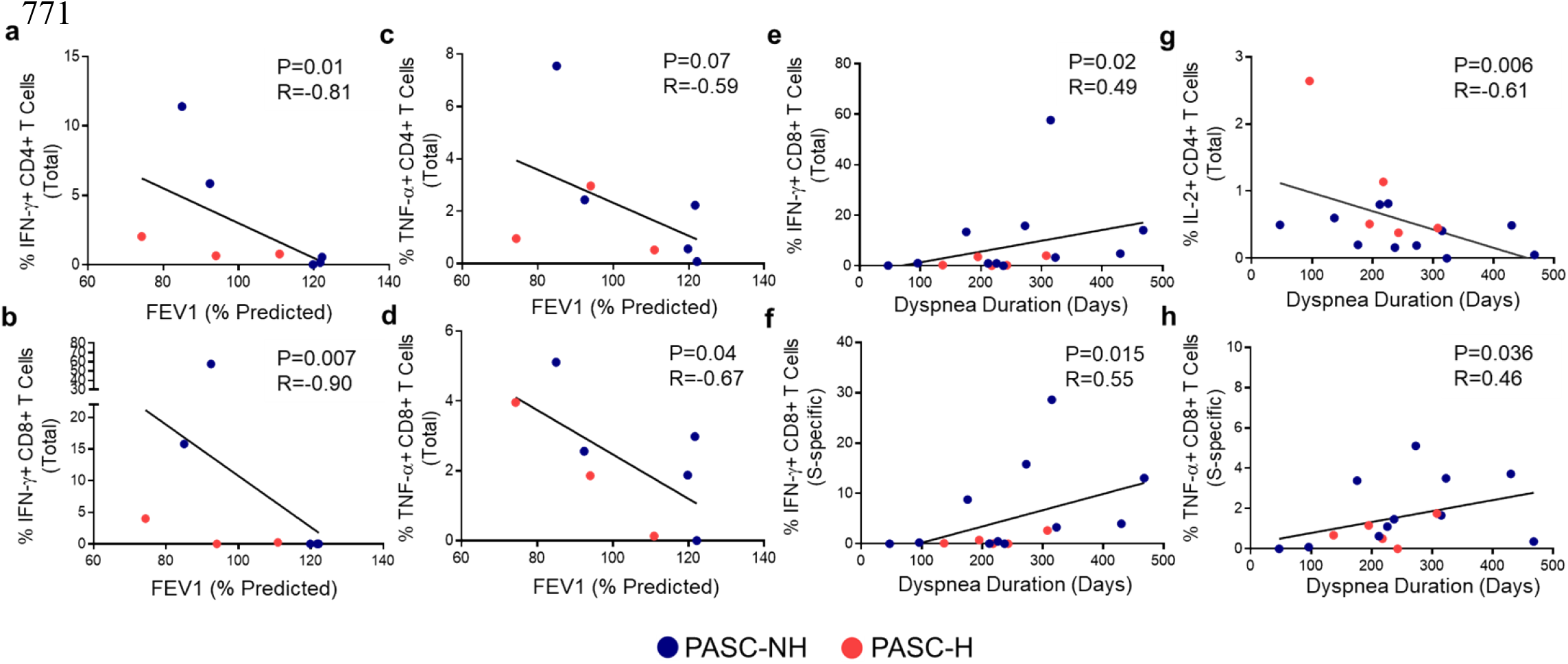
Correlations between SARS-CoV-2 specific T cells and FEVi and symptoms in PASC. Correlations between the total frequency of IFN-γ-producing (a) CD4^+^, (b) CD8^+^ SARS-CoV-2-specific T cells, TNF-α-producing (c) CD4^+^, (d) CD8^+^ SARS-CoV-2-specific T cells, with percent predicted FEV_1_. Correlations between frequencies of (e) total and (f) S-specific IFN-γ-producing CD8^+^ T cells, (g) total IL-2-producing CD4^+^ T cells and (h) S-specific TNF-α-producing CD8^+^ T cells with duration of prolonged dyspnea in days. Each point represents data from one PASC participant. Blue and red symbols represent PASC-NH (not hospitalized) and PASC-Hospitalized participants, respectively. Spearman correlations were used to determine statistical significance.

## Discussion

As the number of SARS-CoV-2 infections accumulate worldwide, PASC is likely to remain a significant health concern for the foreseeable future. We examined SARS-CoV-2-specific immunity in convalescent COVID-19 patients recruited prior to the appearance of the B.1.617.2 “Delta” and B.1.1.529 “Omicron” variants(23). Pulmonary PASC participants with a defined set of prolonged respiratory symptoms had dramatically higher frequencies of SARS-CoV-2-specific T cells in blood compared to participants who had recovered from infection without persistent COVID-19 symptoms. We also found that levels of key plasma inflammatory markers (IL-6 and CRP) were significantly elevated in individuals with ongoing pulmonary PASC and associated with the frequency of SARS-CoV-2-specific T cells. The frequency of SARS-CoV-2-specific T cells in pulmonary PASC participants correlated with reduced lung function and duration of dyspnea, linking the presence of these anti-viral T cells to lung dysfunction. Taken together, these data provide mechanistic insight into the immunopathogenesis of pulmonary PASC.

The most striking feature of PASC is the significantly elevated frequency of SARS-CoV-2-specific TNF-α-producing CD8^+^ T cells. This increased frequency could be detected in response to peptide pools of all the viral structural proteins in comparison to the RC cohort. Interestingly, these T cells were also significantly higher in female PASC participants compared to males, which may contribute to the higher prevalence of PASC in women(1). TNF-α-producing CD8^+^ T cells also expressed the highest levels of Ki-67, indicating recent activation and proliferation. Because the half-life of SARS-CoV-2-specific T cells is between three and five months(24) and most of our PASC participants donated blood over 6 months from symptom onset, it suggests these cells are maintained by viral antigen. The presence of persistent viral reservoirs of SARS-CoV-2 has been proposed as a possible explanation of PASC pathophysiology(27). Studies in macaques and humans demonstrated viral replication can persist months after initial infection in multiple organ systems(28–31) and viral presence in cerebrospinal fluid has been observed in neurological PASC(32). Alternatively, damage resulting from severe disease during acute infection has also been proposed as a cause of PASC(27). However, our results don’t support this idea since sixty percent of our pulmonary PASC cohort initially experienced mild disease(33), yet still developed PASC. Furthermore, there was no difference in the frequency of SARS-CoV-2-specific T cells when hospitalized and non-hospitalized PASC participants were compared. Thus, our findings that pulmonary PASC participants have elevated levels of SARS-CoV-2-specific T cells months after initial infection suggest ongoing viral replication that is maintaining the pool of inflammatory T cells.

The role of T cells in chronic inflammatory conditions is well documented and characterized by the production of TNF-α and other proinflammatory cytokines(34), so we examined the inflammatory markers CRP and IL-6. Both are closely associated with disease severity during acute SARS-CoV-2 infection(35), although previously, no differences in IL-6 levels were found in non-specific PASC when compared to those with resolved infection(36). In contrast, we found that CRP and IL-6 were elevated in pulmonary PASC participants. Levels of CRP in hospitalized PASC participants were significantly higher compared to non-hospitalized PASC participants suggesting prolonged CRP elevation is more strongly associated with initial severity of disease than pulmonary PASC. Elevated IL-6, however, was not different between PASC-H and PASC-NH participants after controlling for sex, age or comorbidities. IL-6 is directly associated with inflammatory lung conditions(37) and targeting IL-6 pathways can effectively treat a variety of inflammatory conditions and decrease mortality in severe COVID-19 cases(38–41). Interestingly, IL-6 levels strongly associated with the frequencies of SARS-CoV-2-specific CD8^+^ T cells which have been shown in other diseases to directly impact tissue-specific monocyte and macrophage production of IL-6 and TNF-α and contribute to feedback loops for innate immune cell recruitment and activation(42, 43) which likely contributes to prolonged respiratory symptoms.

To further understand the role of SARS-CoV-2-specific T cells in pulmonary PASC, we evaluated their associations with lung function. TNF-α impacts asthma progression(44, 45) and chronic obstructive pulmonary disease is associated with IFN-γ-producing T cells(46). In severe COVID-19, decreased pulmonary function is connected to elevated levels of systemic IFN-γ and TNF-α, and analysis of immune cells isolated from bronchoalveolar lavage fluid suggests T cell dysfunction potentially exacerbates tissue damage in severe cases(47–49). These studies indicate a strong connection between T cell cytokine production and lung function, particularly in SARS-CoV-2 infections. However, this association had not been examined in pulmonary PASC. Here, we found that elevated frequencies of IFN-γ- and TNF-α-producing SARS-CoV-2-specific T cells were positively associated with decreased lung function in pulmonary PASC. We also found the duration of dyspnea correlated with increased frequencies of CD8^+^ IFN-γ- and TNF-α-producing SARS-CoV-2-specific T cells and decreased levels of CD4^+^ IL-2 producing T cells. Similar to the effects of systemic cytokines and T cell expression of inflammatory cytokines in other pulmonary conditions, our findings suggest that the presence of persistently activated SARS-CoV-2-specific T cells in PASC likely contributes to lung dysfunction.

Collectively, our findings demonstrate that elevated frequencies of SARS-CoV-2-specific T cells are associated with systemic inflammation and decreased lung function in pulmonary PASC. We observed a striking difference in the frequency of activated and dividing T cells as well as correlations between SARS-CoV-2-specific T cell frequencies and levels of plasma IL-6. Most importantly, we found a strong association between the frequency of SARS-CoV-2-specific T cells and the duration of respiratory symptoms and lung function. While this study examines the responses after infection with one of the early strains of SARS-CoV-2, the characteristics of more recent variants may increase the prevalence of PASC via the same mechanisms supported by our findings(12). Together, these findings suggest pulmonary PASC is in part driven by inflammatory cytokines produced by activated virus-specific T cells, that are likely maintained by persistent virus and contribute to systemic inflammation and prolonged disease morbidity.

## Materials and Methods

### Study participants and sample collection

Adult study participants were recruited from the Denver, Colorado metropolitan area via community flyers, and from the Anschutz Medical Campus Infectious Disease and Pulmonology PASC UCHealth outpatient clinics between July 2020 and April 2021, prior to detection of the Delta or Omicron variants in Colorado(23). Information regarding symptom severity and duration was collected from all participants upon enrollment. 50 mL of blood was collected from study volunteers in sodium heparin tubes (BD, Vacutainer), and plasma and peripheral blood mononuclear cells (PBMCs) were isolated as described previously(50). None of the participants were vaccinated against SARS-CoV-2 prior to sample collection. Participants for this study were only included if they had a documented positive SARS-CoV-2 PCR nasal swab during acute infection and separated into PASC and RC cohorts based on the Center for Disease Control and Prevention definition of PASC(1, 5). We defined pulmonary PASC as having two or more symptoms with a duration longer than 4 weeks from onset or hospital discharge, tussis, or dyspnea present during acute disease and prolonged tussis, dyspnea, and/or fatigue. Demographics of the study population are shown in Table 1 and S1 Table.

### Cytokine and antigen ELISAs

Anti-SARS-CoV-2 Spike S1 IgG and IgA antigens, IL-6 and C-reactive protein (CRP) were assessed using the following ELISA kits per manufacturer protocols: (IgG; Euroimmun - EI2606-9601G, IgA; Euroimmun - EI2606-9601A, IL-6; Invitrogen - 88-7066.22, CRP; Millipore Sigma - RAB0096-1KT). In brief, plasma and standards were diluted per manufacturer’s protocol in sample diluent and, added to pre-coated microplate wells. Following incubation of the wells with biotinylated detection antibody, HRP conjugate, substrate reagent, and stop solution, the plates were read at 450nm.

### T cell stimulation and immunofluorescent staining

The frequency of antigen-specific, cytokine-secreting T cells in blood was determined by intracellular cytokine staining, with minor modifications to our previously published protocol(51). In brief, PBMCs (2-4 × 10^6^ cells) were cultured 5 ml polypropylene tubes in RPMI medium containing 10% human serum and anti-CD28 and anti-CD49d mAbs (each at 1 μg/ml) (S2 Table). Cells were stimulated under the following conditions: peptide arrays of SARS-CoV-2 spike (S) glycoprotein, nucleocapsid (N) protein, membrane (M) protein, (5 μg/ml final concentration of each peptide; BEI Resources from USA-WA1/2020 strain, NR-52402, NR-52404, NR-52403), combined phorbol 12-myristate 13-acetate (PMA) and ionomycin (25 μg/ml and 32.5 μg/ml, respectively; Sigma) or medium alone. S and N arrays were 17- or 13-mer peptides with 10 amino acid overlap, and the M array consisted of 17- or 12-mer peptides with 10 amino acid overlap. Cells were incubated for 6 hours at 37°C in a humidified 5% CO_2_ atmosphere and a 5-degree slant with 1 μg/ml Golgi Plug added after 4 hours. LIVE/DEAD™ Fixable Aqua Dead Cell Stain Kit (Invitrogen L34957) was used per the manufacturer’s protocol after washing. Cells were surfaced stained with the following mAbs: anti-CD3, anti-CD4, anti-CD8, anti-CD27, anti-CD45RA, and anti-PD-1 for 30 min at 4°C. Cells were washed and stored in a fix permeabilization buffer (eBioscience, 421403) overnight at 4°C. Cells were washed in permeabilization buffer and stained with anti-IFN-γ, anti-IL-2, anti-Ki-67, and anti-TNF-α mAbs for 120 min at 4°C, washed, and fixed with 1% formaldehyde. Fluorescence^-1^ (FMO) controls were used in anti-PD-1, anti-CD27 and anti-CD45RA staining. Full information on staining fluorophores are provided in S2 Table.

### Flow cytometry

Cells were analyzed using a LSRII flow cytometer (BD Immunocytometry Systems). At least 1 million events were collected for each tested condition. Antibody capture beads (BD Biosciences) were used to perform electronic compensation. Beads were stained separately with individual mAbs used in the test sample. Data were analyzed using Diva software (BD). Lymphocytes were gated by their forward and side scatter profile. Live and CD3^+^ cells were selected, and expression of CD4 was analyzed in a bivariate dot plot with CD8 to exclude CD4/CD8 double positive T cells. Bi-exponential scaling was used in all dot plots. Expression of CD27, CD45RA, PD-1 and Ki-67 was examined on cytokine-producing cells with at least 100 events to ensure an adequate number of events for analysis(52, 53). FMO controls were used to set gates for determining the percentage of PD-1-expressing T cells. To ensure accuracy and precision of the measurements taken from day-to-day, quality control was performed on the LSRII daily using the Cytometer Setup & Tracking (CS&T) feature of the BD FACSDiva software. This program uses standardized CS&T beads (BD Biosciences) to determine voltages, laser delays, and area scaling to track these settings over time. A manual quality control (QC) using rainbow beads was also performed daily to verify the laser delay and area scaling determined by CS&T.

### Statistics

Statistical analyses were performed using GraphPad-Prism (Graphpad, San Diego, CA). The Mann-Whitney *U* test or Wilcoxon’s matched pairs test were utilized to determine significance of differences between groups. Correlations were calculated using the nonparametric Spearman test. P values of <0.05 were considered statistically significant. To visualize and evaluate differences in expression of multiple cytokines between the PASC and RC cohorts, simplified presentation of incredibly complex evaluations (SPICE) analysis was utilized as well as permutation tests with 10,000 iterations and student T tests for statistical significance where P<0.05 were considered statistically significant. Both of these student T and permutation tests of the SPICE analysis were corrected for 21 concurrent comparisons(54).

### Study approval

This study was approved by the Colorado Multiple Institutional Review Board (COMIRB# 20-1219) at the University of Colorado Anschutz Medical Campus. All participants provided written informed consent prior to any study procedures.

## Acknowledgment

We would like to thank the individuals and organizations who generously shared their time, experience, and materials for the purposes of this project particularly those with PASC who made this possible.

